# Gene sharing networks to automate genome-based prokaryotic viral taxonomy

**DOI:** 10.1101/533240

**Authors:** Ho Bin Jang, Benjamin Bolduc, Olivier Zablocki, Jens H. Kuhn, Simon Roux, Evelien M. Adriaenssens, J. Rodney Brister, Andrew M Kropinski, Mart Krupovic, Dann Turner, Matthew B. Sullivan

## Abstract

Viruses of bacteria and archaea are likely to be critical to all natural, engineered and human ecosystems, and yet their study is hampered by the lack of a universal or scalable taxonomic framework. Here, we introduce vConTACT 2.0, a network-based application to establish prokaryotic virus taxonomy that scales to thousands of uncultivated virus genomes, and integrates confidence scores for all taxonomic predictions. Performance tests using vConTACT 2.0 demonstrate near-identical correspondence to the current official viral taxonomy (>85% genus-rank assignments at 96% accuracy) through an integrated distance-based hierarchical clustering approach. Beyond “known viruses”, we used vConTACT 2.0 to automatically assign 1,364 previously unclassified reference viruses to tentative taxa, and scaled it to modern metagenomic datasets for which the reference network was robust to adding 16,000 viral contigs. Together these efforts provide a systematic reference network and an accurate, scalable taxonomic analysis tool that is critically needed for the research community.

## Main text

Bacteria and archaea modulate the nutrient and energy cycles that drive ocean and soil ecosystems^1–4^, and impact humans by producing metabolites that alter health, behavior, and susceptibility to disease^5^. Viruses that infect these microbes modulate these ‘ecosystem roles’ via killing, metabolic reprogramming and gene transfer^6,7^, with substantial impacts predicted in the oceans^8–10^, soils^11,12^ and human microbiome^13,14^. However, ecosystem-scale understanding is bottlenecked by the lack of universal genes or methods that could facilitate a formalized taxonomy and comparative surveys. In fact, viruses do not share a single gene^15^, and, thus, no analog to microbial 16S rRNA-based phylogenies and OTUs are possible^16^.

Another potential challenge is that some viruses are prone to high rates of gene exchange (i.e., ‘rampant mosaicism’^17^), which, if broadly true, would stymie genome-based prokaryotic virus taxonomy^18^. Fortunately, explorations of viral sequence space are revealing structure^19,20^ and population genetic support for a biological species definition^21^, and new hypotheses to explain variable evolution among prokaryotic viruses^22^. Such findings, alongside rapidly expanding viral genome databases, led the International Committee on Taxonomy of Viruses (ICTV) to present a consensus statement suggesting a shift from the traditional (i.e., phenotypic and genotypic criteria to classify viruses within community-curated taxonomic ranks) approach^23^ towards a genome-centered, and perhaps one-day, largely automated, viral taxonomy^24^. This shift is particularly critical given the modern pace of viral discovery in which, as of March 2018, hundreds of thousands of metagenome-derived viral reference genomes and large genome fragments (369,518 at IMG/VR^25^) now dwarf the 26,223 available from prokaryotic virus sequences in the NCBI GenBank database^26^. Thus, evaluation of approaches to establish a scalable, genome-based viral taxonomy is needed as the implementation of a commonly agreed-upon approach available to the community would be highly desirable.

Multiple genome-based strategies have been proposed to develop such a unified bacterial^15,27–32^, archaeal^33^ and eukaryotic^34^ virus taxonomic framework. For bacterial viruses (“phages”), the first approach targeted phage relationships only by using complete genome pairwise protein sequence comparisons in a phylogenetic framework (the “phage proteomic tree”) and was broadly concordant with ICTV-endorsed virus groupings of the time^15^. Such efforts were not widely adopted, presumably because (i) need was low (few metagenomics studies existed), and (ii) the paradigm was that “rampant mosaicism” would blur taxonomic boundaries and violate the assumptions of the underlying phylogenetic algorithms used in the analyses^17^. Other efforts sought to establish percent of genes shared and percent identity of-shared genes cut-offs to define genera and sub-family affiliations^35,36^, but lacked taxonomic resolution for several virus groups. This lack of resolution was due to the likelihood that the mode and tempo of prokaryotic virus evolution could vary significantly across the viral sequence space^22^. Building upon a prokaryotic classification algorithm, the Genome Blast Distance Phylogeny (GBDP)^37^, a freely accessible online tool (VICTOR) now provides phage genomes for classification via combined phylogenetic and clustering methods from nucleotide and protein sequences^30^. Although a key advance, this method suffers from limited scalability (100-genomes limit) and taxonomic assignment challenges for the many novel, environmental viruses that lack genes shared with reference genomes.

Alternatively, several groups reasoned that the highly variable evolutionary rates across phage sequence space could be examined through gene sharing networks^28,29,38^ to determine whether a meaningful structure, and therefore taxonomic signal, occurs in this space. These networks, based on shared protein clusters (PCs) between viral genomes, were largely concordant with ICTV-endorsed taxa independent of whether monopartite^28^ (a single node type, i.e., viral genomes) or bipartite networks^33,38^ (two node types, i.e., viral genomes and genes) were used. Given these successes, we previously revisited the monopartite gene sharing network approach to establish an iVirus^39^ app (vConTACT) to automate a network-based classification pipeline for prokaryotic virus genomes. Performance tests indicated that the network analytics used by vConTACT produced viral clusters (VCs) that are ~75% concordant with accepted ICTV prokaryotic viral genera, even with seven times more genomes now available^29^. The capacity to incorporate these genomes and accuracy of the network-based analytics have resulted in viral taxonomy applications across large-scale studies of ocean^40,41^, freshwater^42^ and soil^43^, and studies of single-virus amplified genomes (vSAGs)^44,45^. vConTACT 1.0 was an important step forward but could not be used for automatic tentative taxonomic assignments because (*i*) it creates artefactual clusters of both under-sampled genomes (i.e., low number of genomes in a VC) and highly-overlapped regions of sequence space among some genomes^29^, and (*ii*) lacks several key, community-desired features such as confidence metrics for the resultant VCs, a metric for establishing hierarchical taxonomy, and scalability. Here we introduce and evaluate vConTACT v2.0, which updates the network analytics and feature set of the original program. We apply this program to (i) establish a centralized, ‘living’ taxonomic reference network as a foundational community resource and (ii) demonstrate that the updated vConTACT is robust and scalable to modern datasets.

## RESULTS AND DISCUSSION

### vConTACT 2.0 key features and updates

>The underlying goal of vConTACT is to automatically assign viral genomes into relevant established or tentative taxa, with performance assessed relative to ICTV-assigned, manually-curated taxa. Viral reference genomes of a single ICTV genus that are correctly grouped by vConTACT into a single viral cluster (VC) are deemed ‘concordant VCs’. The original vConTACT 1.0 performed well in this area, with ~75% of VCs corresponding to ICTV genera^29^. However, ~25% of VCs did not match ICTV genera (termed ‘discordant VCs’). These mismatches broadly represented three scenarios: (i) VCs that encompass ICTV genera represented by 1-2 genomes (termed ‘undersampled VCs’), (ii) VCs that encompass ICTV genera represented by virus genomes that shared many genes and/or modules with other VCs (termed ‘overlapping VCs’), and (iii) VCs that encompass ICTV genera represented by virus genomes that shared many genes and/or gene modules across genomes within the VC, and within subsets of the genomes in the VC (termed ‘structured VCs’). Further, vConTACT 1.0 lacked several key features to enable broader adoption and utility as described above.

To address these issues and establish vConTACT v2.0, we (i) implemented a new clustering algorithm, (ii) established confidence scores and measures of distance-based taxon separation that are crucial for hierarchical taxonomy, and (iii) optimized expansion to a large-scale viral metagenomic dataset. Briefly, the clustering algorithm was upgraded from Markov cluster (MCL) to ClusterONE^46^ (CL1), resulting in single parameter optimization (i.e., the inflation factor, IF) to determine VC generation being converted to three processes to better disentangle confounding signals across problematic regions of the networks (Online Methods). All three processes consider edge weight, (i.e., degree of connection between genomes), to (i) identify outlier genomes, (ii) detect and separate genomes that bridge overlapping VCs, and (iii) break down structured VCs into concordant VCs through distance-based hierarchical clustering. In addition, to help differentiate between meaningful taxonomic assignments and those that might be artefacts, each VC now receives a topology-based confidence score (value range 0-1), which aggregates information about network topological properties, and a taxonomic (genus) prediction score (value range 0-1), which estimates the likelihood of VCs to be equivalent to a single ICTV genus (Online Methods). In both scores, higher values indicate either more confident linkages (topology-based confidence score) or higher taxonomic agreement (taxonomic prediction score). Therefore, vConTACT 2.0 assigns taxonomy by a two-step clustering approach, in which VCs are first defined using CL1, and then VCs are further subdivided using hierarchical clustering to maximize the taxonomic prediction score. In such cases where VCs were further sub-divided, these are referred to as sub-VCs (benchmarking below).

### Performance comparison of vConTACT versions 1.0 and 2.0

To assess clustering performance of vConTACT v1.0 and v2.0 (hereafter ‘v1.0’ and ‘v2.0’, respectively), we quantified ICTV correspondence from 336 comparisons (Online Methods) against all available ICTV-classified archaeal and bacterial virus genomes (n=2,304, accessed January 2018). Notably, though some combination of family, order, genus and species designations were available for all of these viruses, only 41% (n=940) had genus-level classifications (**Supplementary Table 1**). Our performance comparisons focused on this subset of classified genomes. Composite performance, the sum of six metrics (cluster-wise sensitivity, *Sn*; positive prediction value, *PPV*; geometric accuracy of *Sn* and *PPV*, *Acc*; cluster-wise separation, *Sep_cl_*; complex (ICTV taxon)-wise separation *Sep_co_*; and geometric mean of *Sep_cl_* and *Sep_co_*, *Sep*) was used to assess overall performance of v1.0 and v2.0 (**Fig. 1a**). Each of these metrics has values range from 0 to1 with 1 indicating perfect clustering accuracy and/or coverage (Online Methods). We found that v1.0 organized the 2,304 analysed viral genomes into 305 VCs at its best inflation factor (IF=7), and 77.5% of these were concordant at the genus rank, whereas v2.0 identified 279 VCs, and 79.2% of these were concordant at the genus rank (**Supplementary Table 2**). Moreover, we added to v2.0 a post-processing, Euclidean distance-based hierarchical clustering step to split mismatched VCs. This step accurately and automatically classified 36 additional genera from structured VCs (**Supplementary Table 1**), resulting in the highest composite score of 5.4 (maximum achievable score of 6.0) at the genus rank, with a concordance of 85.0% and accuracy of 96.4%. (**Fig. 1a** and **Supplementary Table 2**). Together, these findings suggest that both upgrading the clustering algorithm and adding hierarchical clustering were critical to improve automatic VC designations.

Next, we assessed how v2.0 handled areas of the reference network that represented discordant VCs. First, 55% of ICTV genera are undersampled (**Supplementary Table 1**), which in a gene-sharing network manifests as weakly connected, small VCs prone to artefactual clustering. In v1.0, undersampled VCs accounted for 64% (28/44) of all discordant VCs, and they could not be resolved by increasing IF values (**Fig. 1b and d** and **Supplementary Table 1**). In contrast, v2.0 automatically and accurately handled these same 28 undersampled VCs (comprising 60 genomes) by splitting the 37 problematic genera into 22 outliers (i.e., genera with only one member) and correctly placing the remaining 38 genomes from 15 genera into 15 VCs (**Fig. 1c and d** and **Supplementary Table 1**). Thus, in instances in which v1.0 performed poorly on undersampled VCs, v2.0 was able to resolve all undersampled VCs into their appropriate ICTV genera.

Second, we evaluated the ability of v2.0 to handle overlapping VCs, which share more genes across VCs than expected, presumably due to gene exchange that could erode structure in the network. In v1.0, overlapping VCs could not be identified. In v2.0 we automated their detection via a ‘match coefficient’ between each VC that measured the connection within-and between-other VCs, and sensitivity analyses established a maximum cluster overlap value of 0.8 as diagnostic (Online Methods). In this way, nine overlapping VCs (ICTV-classified genera only) were detected. These clusters contained 30 viruses across 11 ICTV genera, which included viruses with known mosaic genomes^47^ (e.g., lambdoid or mu-like phages of the *P22virus*, *Lambdavirus*, *N15virus*, and *Bcepmuvirus* genera), temperate phages^48,49^ (i.e., *Mycobacterium* phages of the *Bignuzvirus, Phayoncevirus,* and *Fishburnevirus* genera and *Gordonia* phages of the genus *Wizardvirus*), and three newly-established genera (i.e., *Cd119virus*, *P100virus* and archaeal *Alphapleolipovirus*), all bearing low topology-based confidence scores (averages of 0.29 for these VCs versus 0.50 for concordant VCs; P-value = 2.09e-08, Mann-Whitney U test) (**Supplementary Fig. 1**). Interestingly, this set of viruses within overlapping VCs (74 in total, including non-classified genomes from ICTV) contained 31 phages having a high gene content variation due to extensive gene flow (HGCF, **Fig. 1e**), related to the recently proposed framework of phage evolutionary lifestyles^22^. Further, these VCs contained highly recombinogenic temperate phages, more likely to exchange genes as opposed to low gene content flux (LGCF) phages that follow a predominantly lytic life cycle (**Supplementary Fig. 1b**). Thus, this observation may indicate a high linkage between overlapping genomes and phages with high gene flow. Although unresolvable in v1.0, v2.0 could assign eight of the 11 ICTV genera (24 viruses) into eight ICTV-concordant VCs (**Supplementary Table 1**). The remaining three ICTV genera, all comprised of *Mycobacterium* phages^50^ (six genomes), could not be resolved. This lack of resolution is presumably due to high gene flow resulting from a predominantly temperate lifestyle that is associated with an exceptionally high fraction (avg = 69%) of genes shared across VCs (**Supplementary Table 3**). Undoubtedly, these genomes are the most challenging to classify, and may not be amenable to automated taxonomy. Whether such highly recombinogenic genomes are the exception or the norm across environments is unknown.

Third, structured VCs contained genomes that our gene sharing networks placed into a single VC (due to many shared genes and/or gene modules across all the member genomes), whereas ICTV delineated multiple genera (due to subsets of the genomes also sharing additional genes). V1.0 qualitatively and selectively handled these structured VCs via decomposing hierarchical patterns of gene sharing^27^. In v2.0, we formalized an optimized, quantitative hierarchical decomposition distance measure (9.0, Online Methods, **Fig. 2c**, and **Supplementary Fig. 2**) that maximized composite scores of two geometric mean values of performance metrics (*Acc* and *Sep*; Online Methods) that divide discordant VCs into concordant (to ICTV genera) sub-VCs, and used this distance as a generalized threshold. In the v2.0 network, 31 discordant VCs contained 101 phage and two archaeal virus genera, in which 23 (74%) were structured VCs spanning 86 genera (**Fig. 2a,b** and **Supplementary Table 1**). This v2.0 approach resolved 30% (26 of 86) of these ICTV genera from 6 of the 23 structured VCs (**Fig. 2c**). Curiously, one such structured VC was comprised of T4-like phages (of which nine out of ten T4-related genera were resolved; **Supplementary Note 1**), in which hierarchical ‘T4 core’ and ‘cyano T4 core’ gene sets are well documented^51^. In our networks, the T4-like phages represent a single VC, but with sub-VCs that are consistent with ICTV-established genera (VC 1 in **Fig. 2c** and **Supplementary Table 1**). Extrapolating from this network, we interpret structured VCs to represent areas of viral sequence space that are well-sampled to the point that the core gene sets that define a virus (capsid, tail, replication machinery) establish the VC in the network, whereas ecologically diverse viral genomes within the VC reveal structure due to niche-defining genes that represent adaptation to diverse environments and/or hosts. We posit that the 19 structured VCs that cannot be resolved towards ICTV concordance (**Fig. 2c** and **Supplementary Table 1**), represent either regions of the network where niche-defining genomic information is lacking or may require complementary phenotypic or evolutionary evidence to establish ICTV genera, as done for the archaeal fuselloviruses (VC42) and bacterial microviruses (VCs 30 and 49). Thus, whether these structured VCs result from lack of resolution in v2.0 or from genera needing ICTV revision remains an outstanding question.

Finally, given such strong performance, we suggest that this gene sharing network already offers significant new taxonomic insights. First, as described earlier, only 41% of the 2,304 reference virus genomes are classified by ICTV at the genus rank. Thus, we propose that the remaining 1,364 currently genus-unclassified reference viruses, which organized into 304 well-supported hierarchically decomposed sub-VCs (**Supplementary Table 1**), represent genomes from *bona fide* novel virus genera. This finding, if officialised, immediately doubles established viral taxonomy and invites a framework for manual curation of these automatic assignments, which in itself will improve future vConTACT analytic performance. As first evidence of the value of such an iterative process, we note that v2.0 clustering suggested an alternative taxonomy among ten current ICTV genera: *Barnyardvirus*, *Bcep78virus*, *Bpp1virus*, *Che8virus*, *Jerseyvirus*, *P68virus*, *Pbunavirus*, *Phietavirus*, *Phikmvvirus*, and *Yuavirus* (**Supplementary Fig. 3** and **Note 2**), and manual inspection had already recommended some of these ICTV genera be revised (e.g. *Phikmvvirus* viruses, ICTV proposal 2015.007a-Db). An automated vConTACT-based approach would systematically identify such problematic taxa and drastically speed up these critical revisions as new data become available.

### vConTACT v2.0 is scalable to modern virome datasets

A major bottleneck regarding automated taxonomic assignments is the ability to robustly integrate large sets of newly discovered virus genomes. To evaluate this concern, we added ~16K curated viral genomes and large genome fragments from the Global Ocean Virome (GOV) dataset^40^ to our reference network. We added these genomes and genome fragments in successive 10% increments (i.e., 0%-10%,[…], 0%-100%), to assess the impact of various data scales on the reference network stability of VC assignments. Network changes were tracked by assessing (i) network performance metrics (*Sn*, *Acc* & *PPV,* as above), (ii) ‘normalized mutual information’(NMI), as a measure of VC similarity (values range from 0 to 1 with 0 indicating that none of the original member genomes within a VC remained in that same VC and 1 indicating that all members in a VC remain in that same VC across time), and (iii) ‘change centrality’ (CC), reflecting how much each node’s connections changed as more sequences were added to the network (values range from 0-1 with 0 indicating no change and 1 indicating complete change), classified over three ‘change intensity’ groups: low (0 - 0.283), medium (0.283 - 0.506) and high (0.506 - 0.999) groups (Online Methods). Although CC indicates changes in connections between nodes, these may still remain in a given VC, albeit re-shuffled. Together, NMI and CC assess the impact of additional data on the network clusters and topology, respectively, while *Sn*, *Acc* and *PPV* assess concordance with ICTV taxonomy.

All measures indicated that most network changes occurred with early additions of the novel GOV data (up to 20-30% of the dataset), with the network largely stabilized after that (**Fig. 3**). For example, *Acc* (mean value of *Sn* and *PPV*) is reduced by 12% when only using 20% of the GOV data, but stabilizes at a ~7% decrease (**Supplementary Fig. 4**); similar responses were observed in NMI (**Fig. 3b**). This initial drop appears driven by formation of novel, undersampled VCs, a disruptive effect similarly observed with undersampled ICTV genera bearing low quality or confidence in VC membership. With more data, undersampled VCs reach ‘saturation’, which increases confidence scores for these new VCs and buffers from further disruption. This stabilization is likely due to strong intra-cluster forces (within VCs) vastly out-weighing inter-cluster forces (between VCs). The lasting minimal decrease represents the novelty of sequence space in GOV relative to RefSeq and the fact that these additions are commonly large genome fragments rather than complete genomes. Sequential CC analysis showed minimal impact on the RefSeq network structure and VC membership, as 85% of reference genomes had low-to-medium change, whereas 0.05% of genomes experienced high change. The remaining 15% were classified as either singleton, outlier, or overlaps. These data support a similar pattern as NMI fluctuations (**Fig. 3d**): as data accumulated, fewer and fewer nodes or VCs were impacted due to new data influencing only pre-existing areas in the network. Therefore, as a network grows in scale, adding new data mostly similar to pre-existing data will have minimal impact on the underlying network structure (e.g., adding new marine data to a marine network), as newly added data is already “represented,” whereas utterly novel data will generate novel VCs and increase CC values. Indeed, most unaffected VCs (CC = 0) were non-marine or soil in origin e.g., *Andromedavirus* viruses, *Saetivirus* viruses, two archaeal viruses (Methanobacterium virus psiM2, Methanothermobacter virus psiM100), *Thermus* phages, or cyanobacterial mat viruses.

As contigs accumulate, the number of VCs also increases linearly (R^2^ = 0.998, P-value = 1.2 x 10^-12^). We examined whether GOV data may partially resolve ICTV outlier and singleton genomes. More data should create new connections to singletons, whereas outliers may get connected to new or existing VCs. Out of 38 single-member VCs of singleton and outlier genomes (**Supplementary Fig. 5**), three *Mycobacterium* phages clusters were improved, with two other *Mycobacterium* viruses genomes merged into six-genera heterogeneous VCs. Together, this analysis suggests that v2.0’s underlying methodology is sufficiently robust to handle large amounts of data. With 100% of GOV added (16,960 total contigs), 919 new VCs are created, representing potentially 919 new viral genera over existing RefSeq genomes.

### Community availability and future needs

The utility of v2.0 depends upon its expert evaluation and community availability. To maximize this evaluation, members of the ICTV Bacterial and Archaeal Viruses Subcommittee were invited as co-authors to critique the work, and we made the resulting optimized tool available in two ways. First, the source code is available through Bitbucket (https://bitbucket.org/MAVERICLab/vcontact2 as a downloadable python package. Second, v2.0 is available as an app through iVirus^39^, the viral ecology apps and data resource embedded in the CyVerse Cyberinfrastructure, with detailed usage protocols available through Protocol Exchange (https://www.nature.com/protocolexchange/) and protocols.io (https://www.protocols.io/). Finally, the curated reference network is available at each of these sites.

Although v2.0 performance metrics are strong and provide a critically needed, systematic reference viral taxonomic network, limitations still remain. First, our reference network needs to be rebuilt each time new data are added. Avoiding this reconstruction step will require the development of approximation methods and/or a placement algorithm (akin to PPlacer for 16S phylogenies^52^) to incorporate new data. Second, although v2.0 handles reference prokaryotic virus genomes (including ssDNA or dsDNA phages) and large GOV genome fragments, this framework has not been designed, tested or validated for eukaryotic viruses, which pose unique computational challenges^34^. Third, shorter prokaryotic virus genomes and genome fragments (e.g., ≤ 3 PCs or ≤ 5 genes) are of low statistical power in the v2.0 framework, and will require new solutions to establish higher confidence VCs. Fourth, genomes identified as singletons, outliers or overlapping are currently excluded from the gene-sharing network. Although singletons and outliers can be resolved by the addition of new data, overlapping VCs can remain challenging to resolve, particularly for the HGCF phages^22^ that are highly recombinogenic. Such rampantly mosaic virus genomes are problematic for viral taxonomy. However, they are identifiable in the networks and, at least to date, represent the minority of known viral sequence space. Most (~75%) are LGCF viruses that remain amenable to automated genome-based viral taxonomy. Whether this situation will remain so awaits further exploration of viral sequence space—particularly where temperate phages may predominate (e.g., soils^53^, human gut^54^). For now, we propose vConTACT 2.0 as a tool that offers a robust, systematic and automatic means to aid the classification of bacterial and archaeal viruses.

## METHODS

### Data sets

Full-length viral genomes were obtained from the National Center for Biotechnology Information (NCBI) viral reference dataset^26,55^ (‘ViralRefSeq’, version 85, as of January, 2018), downloaded from NCBI's viral genome page (https://www.ncbi.nlm.nih.gov/genome/viruses/) and eukaryotic viruses were removed. The resulting file contained a total of 2,304 RefSeq viral genomes including 2,213 bacterial viruses and 91 archaeal viruses (**Supplementary Table 1**). In parallel, the ICTV taxonomy (ICTV Master Species List v1.3, as of February, 2018) was retrieved from the ICTV homepage (https://talk.ictvonline.org/files/master-species-lists/). ICTV-classifications were available for a subset of genomes at each taxonomic rank, and final dataset included; 884 viruses from two orders, 974 viruses from 23 families, 363 viruses from 28 subfamilies, and 975 viruses from 264 genera. To maintain hierarchical ranks of taxonomy, we manually incorporated 2016 and 2017 ICTV updates^56–58^ to NCBI taxonomy when ICTV taxonomy was absent.

### Network construction

A total of 231,165 protein sequences were extracted from the 2,304 viral genomes (above). To group protein sequences into homologous protein clusters (PCs)^29^, all proteins were subjected to all-to-all BLASTP^59^ searches (default parameters, cut-offs of 1E^-5^ on e-value and 50 on bit score). A subsequent application of the MCL with inflation factor 2.0 grouped 204,540 protein sequences into 25,510 PCs, with the remaining 26,625 proteins being to singletons (those that do not have close relatives). The resulting output was parsed in the form of a matrix comprised of genomes and PCs (i.e., 2,304 × 25,510 matrix). We then determined the similarities between genomes by calculating the probability of finding a common number of PCs between each pair of genomes, based on the following hypergeometric equation as per Lima-Mendez et al^28^:

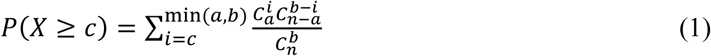

in which *c* is the number of PCs in common; *a* and *b* are the numbers of PCs and singletons in genomes A and B, respectively; and *n* is the total number of PCs and singletons in the dataset. A score of similarity between genomes was obtained by taking the negative logarithm (base 10) of the hypergeometric P-value multiplied by the total number of pairwise genome comparisons (i.e., *2,304* × *2,303*). Genome pairs with a similarity score ≥1 were previously shown to be significantly similar through permutation test of PCs and/or singletons between genomes^29^. Afterwards, a gene (protein)-sharing network was constructed, in which nodes are genomes and edges connect significantly similar genomes. This network was visualized with Cytoscape software (version 3.6.0; http://cytoscape.org/), using an edge-weighted spring embedded model, which places the genomes sharing more PCs closer to each other.

### Parameter optimization of vConTACT v1.0 and 2.0

Due to different criteria for parameter optimization between the clustering methods, different number and size of the clusters are often generated, which can make objective performance comparisons difficult^60^. Thus, to more comprehensively compare performance, v 1.0’s MCL-based VCs were generated at inflation factors (IFs) of 2.0 to 7.0 by 1.0 increments, with an optimal IF of 1.4 showing the highest intra-cluster clustering coefficient (ICCC)^28^ (**Supplementary Table 1** and **Supplementary Fig. 6**). CL1, which was incorporated into a new version of vConTACT (v2.0), operates in multiple stages of complex detection^46^. Unlike the MCL that uses a single parameter^28^, CL1 uses a set of parameters, which can act as the threshold for each stage of complex detection. For example, as four main parameters of CL1, the minimum density, node penalty, the haircut, and the overlap automatically quantifies (i) the cohesiveness of cluster, (ii) the boundaries of the clusters (outliers), and (iii) the size of overlap between clusters, respectively^46^. Of these parameters, the first two are used to detect the coherent groups of VCs as follows:

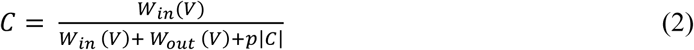

in which *W_in_(V)* and *W_out_(V)* are the total weight of edges that lie within cluster *V* and that connect the cluster *V* and the rest of the network, respectively, |*C*| is the size of the cluster, *p* is a penalty that counts the possibility of uncharted connections for each node.

As another parameter of CL1, the haircut can find loosely connected regions of the network (outliers) by measuring the ratio of connectivity of the node *g* within the cluster *c* to that of its neighbouring node *h* as:

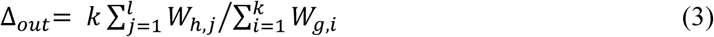

in which *k* is the number of edges of the node *g*, and *W* is the total weight of edges of the respective nodes *g* and *h*. If the total weight of edges from a node (*h*) to the rest of the cluster (*c*) is less than x times that we specified the average weight of nodes (*g*) within the given cluster, CL1 will remove the node (*h*) from a given VC and place it into the outlier.

Additionally, CL1 can specify the maximum allowed overlap (*ω*) between two clusters, measured by the match coefficient, as follow:

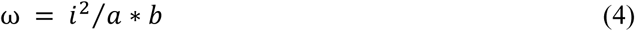

in which *i* is the size of overlap, which is divided by the product of the sizes of the two clusters under consideration (*a* and *b*). Since CL1 identifies overlap between VCs, it can consequently find both hierarchical and overlapping structures of viral groups. This capability is a significant improvement over v1.0, given v1.0’s MCL cannot handle modules with overlaps^7^. Specifically, CL1 (i) finds cluster(s) having less than maximum value of specified overlap threshold (above) and (ii) merges these clusters together with their interacting cluster(s) to make the results easier to interpret. Thus, in the resulting output file, viral groups (or clusters) having the identical member viruses can be found in multiple clusters, called ‘overlapping clusters’ (**Supplementary Table 1**). CL1 was run with varying conditions for these four parameters (minimum density ranging from 0 to 1 by 0.1 increments; node penalty from 1 to 10 by 1.0; haircut from 0 to 1 by 0.05; overlap from 0 to 1 by 0.05) and default settings for other parameters: 2 as minimum cluster size, weighted as edge weight, single-pass as merging, unused nodes as seeding. We therefore obtained a total number of 53,361 clustering results, which we evaluated individually to yield the highest performance on taxonomic data set (above), in terms of geometric mean value of prediction accuracy (*Acc*) and clustering-wise separation (*Sep*, see next section), as previously described^61^ We then used minimum density = 0.3, node penalty = 2.0, haircut = 0.65, and overlap = 0.8 to derive the final set of clusters, resulting in a total of 279 VCs (**Supplementary Table 1**). As a post-clustering step of v2.0, all VCs including discordant clusters (those comprising ≥ 2 taxa) were further hierarchically separated into sub-clusters using the unweighted pair group method with arithmetic mean (UPGMA) with pairwise Euclidean distances implemented in Scipy. To optimize the distance-based sub-clustering of VCs, we assessed the distances of sub-clusters across all the VCs. These distances (ranging from 1 to 20 in 0.5 increments) maximized the geometrical mean values of the prediction accuracy (*Acc*) and clustering-wise separation (*Sep*) at the ICTV genus rank (see next section). This optimization resulted in the distance of 9.0 yielding the highest composite score of *Acc* and *Sep* (**Supplementary Fig. 2**). Notably, vConTACT v2.0 was designed to help users optimize (i) parameters for grouping of genomes/contigs into VCs and (ii) distance for post-decomposition of VCs into sub-clusters. This tool automatically evaluates the robustness of VCs and sub-clusters, respectively, based on the external performance evaluation statistics (below).

### Performance comparison between vConTACT v1.0 and v2.0

Since the external measures such as precision, recall, and others often neglect overlapping clusters, which might not reflect the true performance of CL1, we used 6 external quality metrics that were successfully used for performance comparison between MCL and CL1^61^ (see below). Specifically, the performance of v1.0 (MCL) and v2.0 (CL1 alone and CL1 + hierarchical sub-clustering, respectively) were evaluated based on : (i) cluster-wise sensitivity, *Sn* (ii) positive predictive value, *PPV* (iii) geometric accuracy of *Sn* and *PPV*, *Acc* (iv) cluster-wise separation, *Sep_cl_* (v) complex (ICTV taxon)-wise separation *Sep_co_*, and (vi) geometric mean of *Sep_cl_* and *Sep_co_*, *Sep*. As an internal parameter, we computed the intra- and inter-cluster proteome similarities (fraction of shared genes between genome that are within the same VCs and different VCs, respectively). For vConTACT v1.0, clustering result yielding the highest clustering accuracy value (inflation of 7.0) was subsequently used for comparison to v2.0’s clusters and sub-clusters.

To generate six external measures, we first built a contingency table *T*, in which row *i* corresponds to the *i^th^* annotated reference complex (i.e., ICTV-recognized order, family, subfamily, or genus), and column *j* corresponds to the *j^th^* predicted complex (i.e., sub-/clusters). The value of a cell *T_ij_* denotes the number of member viruses in common between the *i^th^* reference complex and *j^th^* predicted complex. Here, *N_i_* is the number of member viruses belonging to reference complex *n*. *Sn* and *PPV* are then defined as follows:

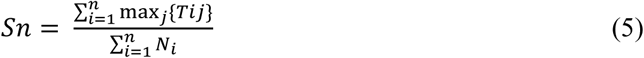

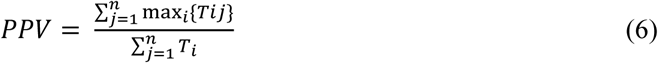

Generally, higher *Sn* values indicate a better coverage of the member viruses in the real complexes, whereas higher *PPV* values indicates that the predicted clusters are likely to be true positives. As a summary metric, the *Acc* can be obtained by computing the geometrical mean of the *Sn* and *PPV* values:

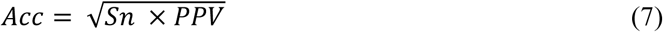

With the same contingency table used for *Sn*, *PPV*, and *Acc*, we calculated the averages of complex-wise separation *Sep_co_*, and cluster-wise separation *Sep_cl_*, respectively, below:

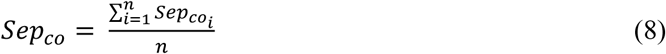

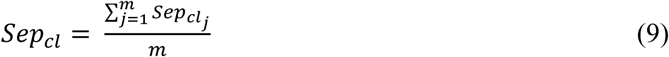

High *Sep_co_* and *Sep_cl_*, (both have maximal values of 1.0) indicate how well a given complex is isolated from the other complexes and a cluster from other clusters, respectively. To estimate these separation results as a whole, the geometric mean (clustering-wise separation; *Sep*) of *Sep_co_* and *Sep_cl_* was computed:

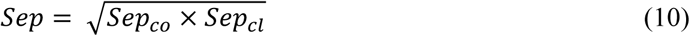

High clustering-wise separation values indicate a bidirectional correspondence between a sub-/cluster and each ICTV taxon: maximal value of 1.0 can be obtained when a sub-/cluster corresponds perfectly to each taxon.

As an internal measure, the fraction of PCs^29^ between two genomes (i.e., proteome similarity) was computed by using the geometric index (G). The proteome similarity was estimated as:

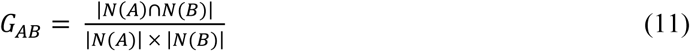

in which *N(A)* and *N(B)* indicate the number of PCs in the genomes of A and B, respectively. A total of 400,234 pairs of genomes with >1% proteome similarity are shown in **Supplementary Table 3**.

### Clustering-based confidence score

To generate the confidence score per sub-cluster, we used four confidence scoring methods, as previously described^62,63^, with some modifications. Three of them exploit the network topology properties by assessing (i) the significance of clustering coefficient, (ii) the weight of cluster quality, and (iii) the probability of cluster quality. We then used combined these three values into an aggregate topology-based confidence score.

Specifically, for the significance of the clustering coefficient, we quantified the fidelity (*F*) of the edge (*p*) by calculating cumulative hypergeometric P-values using Equation 1 (above) between sub-clusters. The fidelity values are lower (close to 0) for the genomes having the higher number of shared genes. We then defined the confidence of sub-cluster cohesiveness as the product of the fidelity values of total edges (i.e., *p1 and p2*) within the sub-cluster *c* as below:

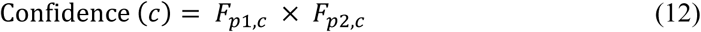

For the second scoring method, we computed the quality (*Q*) of sub-cluster (*c*) as:

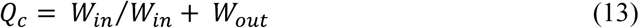

in which *W_in_* and *W_out_* are the total weight of edges that lie within sub-cluster *c* and across others, respectively. For the third method, we evaluated the P-value of a one-sided Mann-Whitney U test for in-weights and out-weights of sub-clusters. The rationale behind this test is that sub-clusters with a lower P-value contains significantly higher in-weights than out-weights, thus indicative that a formed sub-cluster is valid, and not a random fluctuation. All pairs of three values above were then incorporated into the topology-based confidence score with the Spearman rank correlation coefficient by using in-house python scripts and Scipy. Along with this confidence score, we quantified the likelihood that each sub-cluster corresponds to an ICTV-sanctioned genus (or equivalent) by using distance threshold that are specified at the ICTV genus rank, which we refer to as “taxon predictive score”. This score can be calculated as:

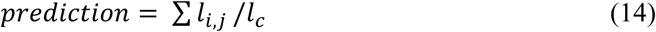

Specifically, for a sub-cluster (*c*) having the genus-level assignment, vConTACT v2.0 automatically measures the maximum distance between taxonomically-known member viruses and calculate the scores by dividing the sum of links having less than the given maximum distance threshold between nodes (*i* and *j*) by the total number of links (*l_c_*) between all nodes. For a sub-cluster that does not have the genus-level assignment, v2.0 uses Euclidean distance of 9.0 that can maximize the prediction accuracy and clustering-wise separation (see above) as distance threshold.

### Measuring effect of GOV on network structural changes

GOV contigs (14,656) were added in 10% increments (randomly selected at each iteration) to NCBI Viral RefSeq and processed using vConTACT 2.0 with one difference – Diamond^64^ instead of BLASTp was used to construct the all-versus-all protein comparison underlying the PC generation. Once generated, vConTACT 2.0 networks were post-processed using a combination of the Scipy^65^, Numpy, Pandas^66^ and Scikit-learn^67^ python 3.6 packages. Networks were rendered using iGraph^68^. To calculate NMI, each network’s genomes and their VC membership was compared in pairwise fashion to all other networks using the “adjusted mutual info score” function of Scikit-learn. Intra-cluster distances were calculated using the agglomerative clustering functions “linkage” with distance calculated from shared PCs using the cluster average (also known as UPGMA), and novel clusters identified using the “fcluster” function of Scipy’s hierarchical clustering. In parallel, the method to calculate change centrality was calculated as described previously^69^. CCs were calculated in a successive way, in which each addition was compared to Viral RefSeq 85 independently of other additions (0% versus 10%, 0% vs 20%, […], 0% vs 100%).

### Code availability

The vConTACT v2.0 package is freely distributed through Bit Bucket as a python package (https://bitbucket.org/MAVERICLab/vcontact2).

## ACKNOWLEDGEMENTS

We thank Laura Bollinger, Gareth Trubl, and Igor Tolstoy for their comments on improving the manuscript, as well as Wesley Zhi-Qiang You for helping push the network analytics. High performance computational support was provided as an award from the Ohio Supercomputer Center to MBS. Funding was provided in part by the Department of Energy’s Genome Sciences Program Soil Microbiome Scientific Focus Area award (#SCW1632) to Lawrence Livermore National Laboratory; an NSF Biological Oceanography award (OCE#1536989), and a Gordon and Betty Moore Foundation Investigator Award (#3790) to MBS. Funding was provided to JRB by the Intramural Research Program of the NIH, National Library of Medicine. The work conducted by the U.S. Department of Energy Joint Genome Institute is supported by the Office of Science of the U.S. Department of Energy under Contract DE-AC02-05CH11231 to SR. This work was funded in part through Battelle Memorial Institute’s prime contract with the US National Institute of Allergy and Infectious Diseases (NIAID) under Contract No. HHSN272200700016I to JHK. The content of this publication does not necessarily reflect the views or policies of the US Department of Health and Human Services or of the institutions and companies affiliated with the authors.

## AUTHOR CONTRIBUTIONS

HBJ, BB and MBS designed the study. OZ and MBS wrote the manuscript with significant contributions from all co-authors. HBJ and BB performed the statistical and network analyses.

## COMPETING INTERESTS

The authors declare no competing interests.

## MATERIALS & CORRESPONDENCE

Correspondence and material requests should be addressed to Matthew B. Sullivan at.

---

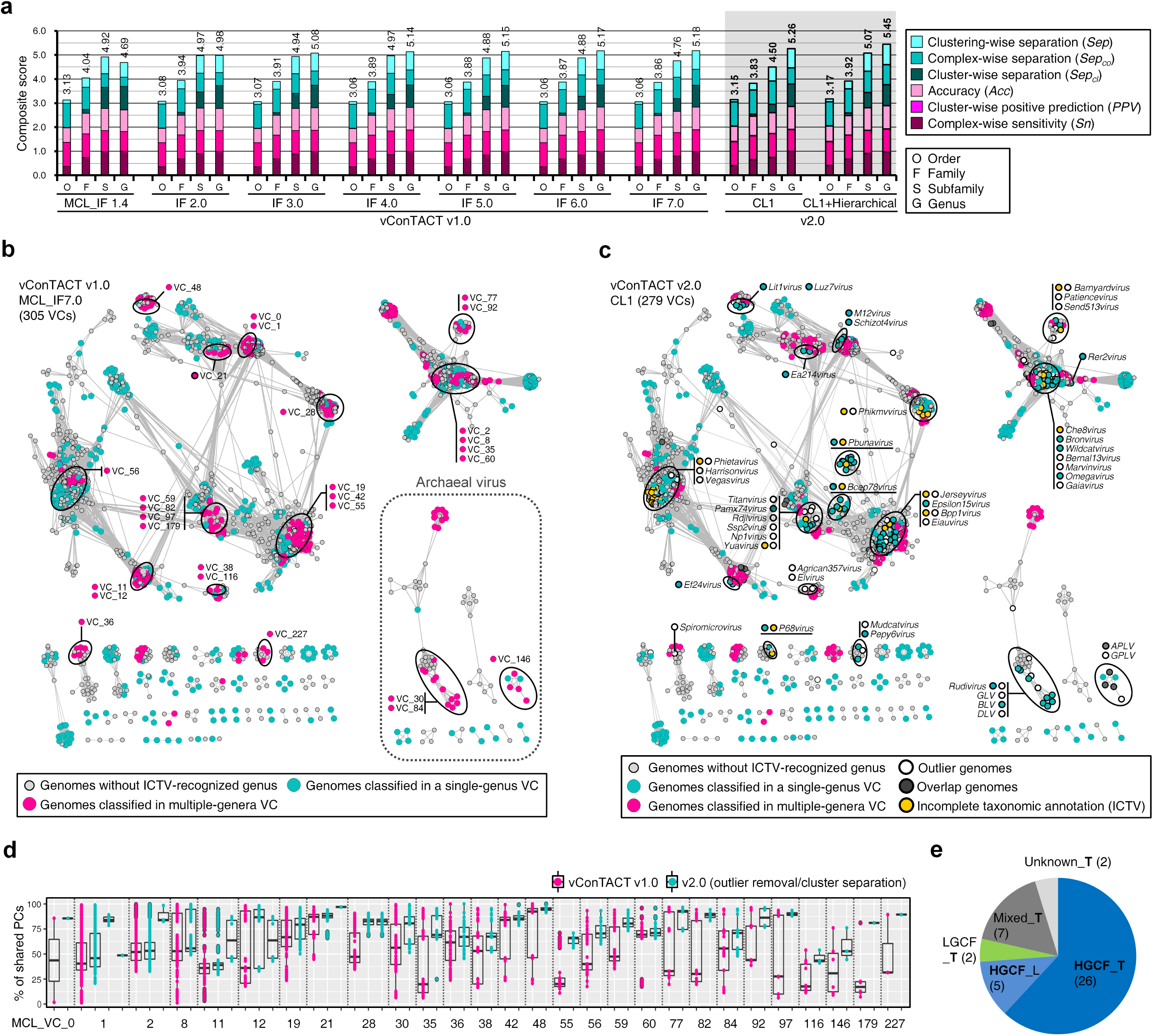

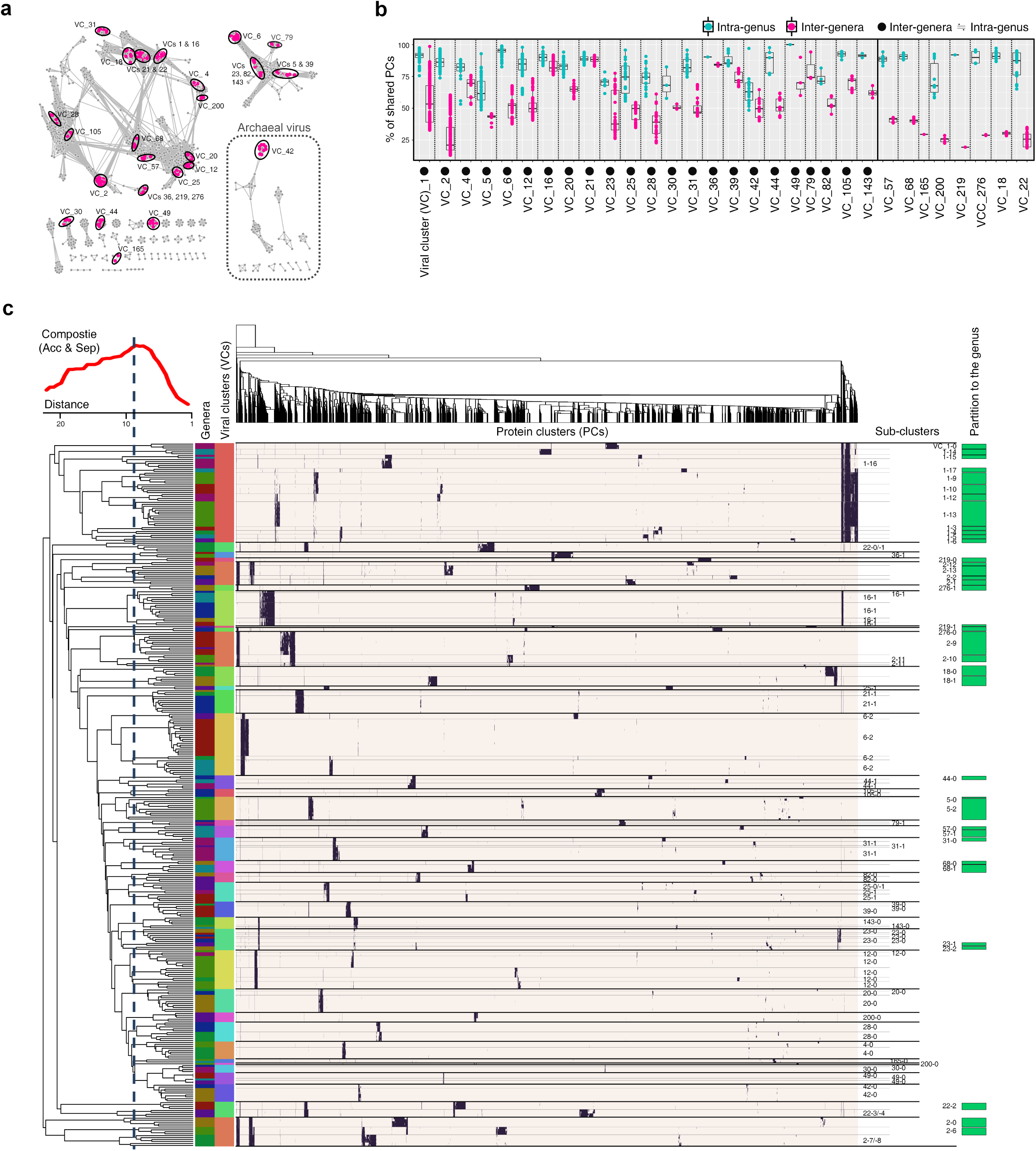

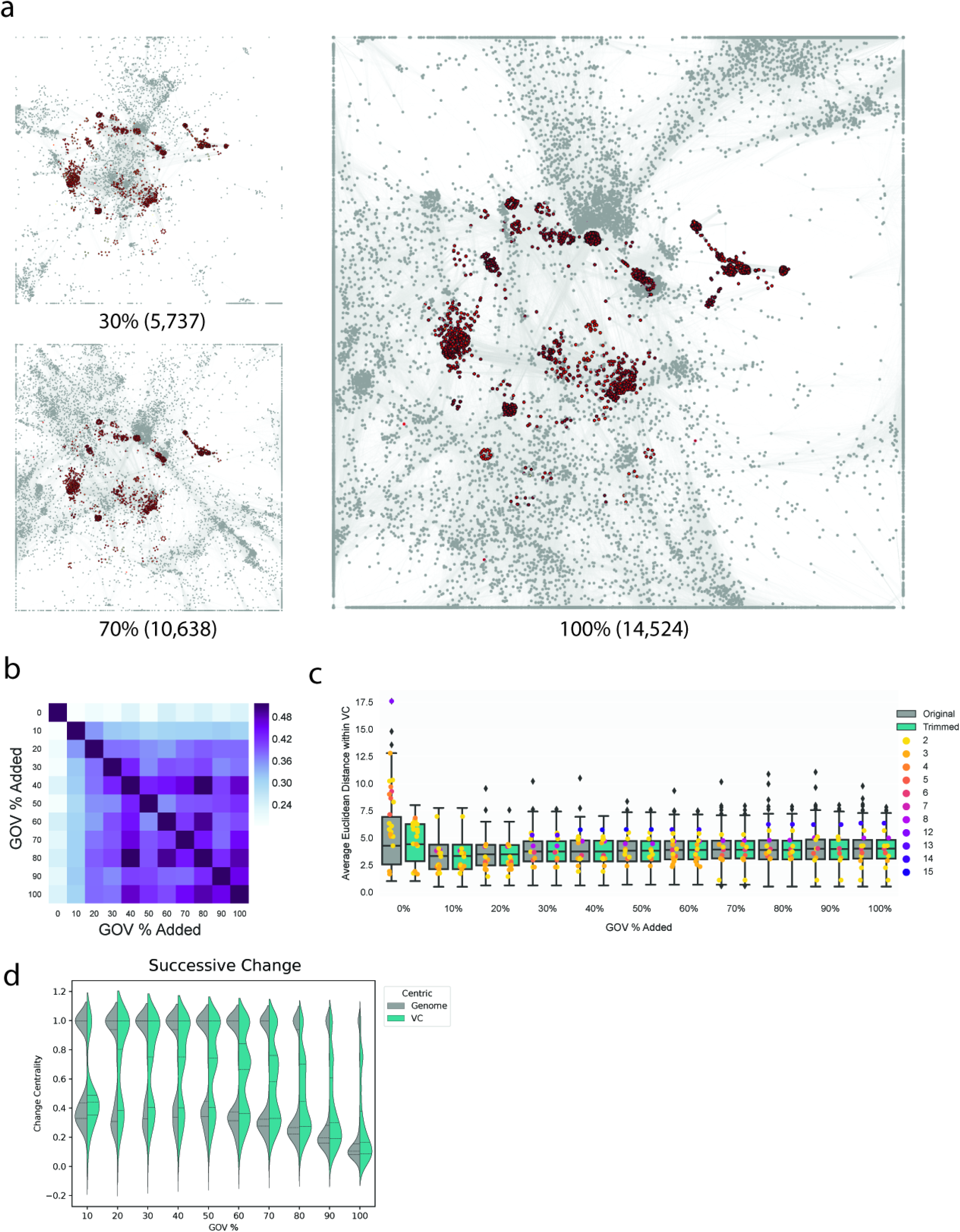

