## Supplementary figures for "Gene sharing networks to automate genome-based prokaryotic viral taxonomy"

a

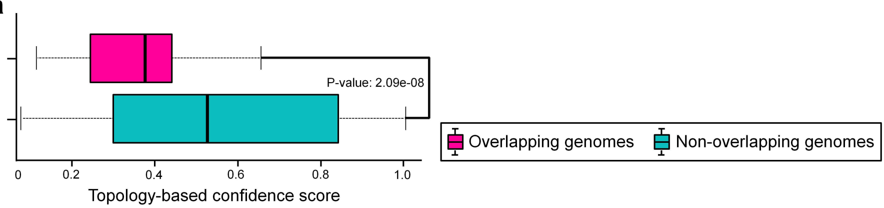


b

Evolutionary mode

HGCF LGCF Mixed Unknown Uncharacterized

| Name | Accession | ICTV-Genus | vConTACT_v2.0_VCs | Temperate (empirical) | Temperate (bioinformatically predicted) | Evolutionary mode |
|---|---|---|---|---|---|---|
| Enterobacteria_phage_HK629 | NC_019711 | Lambdavirus | 13/218 | yes | yes | HGCF |
| Escherichia_virus_Lambda | NC_001416 | Lambdavirus | 13/218 | yes | yes | HGCF |
| Enterobacterial_phage_mEp213 | NC_019720 | | 13/218 | yes | yes | HGCF |
| Enterobacteria_phage_mEp043_c-1 | NC_019706 | | 13/218 | yes | yes | HGCF |
| Enterobacteria_phage_mEp460 | NC_019716 | | 218/279 | yes | yes | HGCF |
| Enterobacteria_phage_phi80 | NC_021190 | | 218/279 | yes | yes | HGCF |
| Enterobacteria_phage_cdtI | NC_009514 | | 218/279 | yes | yes | HGCF |
| Lactococcus_phage_phiLC3 | NC_005822 | | 85/113 | yes | yes | HGCF |
| Lactococcus_phage_r1t | NC_004302 | | 85/113 | yes | yes | HGCF |
| Lactococcus_phage_TP901-1 | NC_002747 | | 85/277 | yes | yes | HGCF |
| Lactococcus_phage_Tuc2009 | NC_002703 | | 85/277 | yes | yes | HGCF |
| Salmonella_enterica_bacteriophage_SE1 | NC_011802 | | 13/32 | yes | yes | HGCF |
| Salmonella_phage_epsilon34 | NC_011976 | | 13/32 | yes | yes | HGCF |
| Salmonella_phage_SPN9CC | NC_017985 | | 13/32 | yes | yes | HGCF |
| Salmonella_phage_ST160 | NC_014900 | | 13/32 | yes | yes | HGCF |
| Salmonella_phage_vB_SemP_Emek | NC_018275 | | 13/32 | yes | yes | HGCF |
| Enterobacteria_phage_ST104 | NC_005841 | | 13/32 | yes | yes | HGCF |
| Salmonella_phage_ST64T | NC_004348 | P22virus | 13/32 | yes | yes | HGCF |
| Salmonella_virus_HK620 | NC_002730 | P22virus | 13/32 | yes | yes | HGCF |
| Salmonella_virus_P22 | NC_002371 | P22virus | 13/32 | yes | yes | HGCF |
| Shigella_virus_Sf6 | NC_005344 | P22virus | 13/32 | yes | yes | HGCF |
| Streptococcus_phage_315.4 | NC_004587 | | 192/228 | yes | yes | HGCF |
| Temperate_phage_phiNIH1.1 | NC_003157 | | 192/228 | yes | yes | HGCF |
| Clostridium_phage_phiCD505 | NC_028764 | | 45/256 | yes | yes | HGCF |
| Clostridium_phage_phiMMP02 | NC_019421 | | 45/256 | yes | yes | HGCF |
| Clostridium_virus_phiCD27 | NC_011398 | Cd119virus | 45/256 | Uncharacterized | yes | HGCF |
| Enterobacteria_phage_Sf101 | NC_027398 | | 13/32 | Uncharacterized | yes | HGCF |
| Salmonella_phage_SEN22 | NC_028696 | | 13/32 | Uncharacterized | yes | HGCF |
| Listeria_phage_List-36 | NC_024364 | P100virus | 2/280 | Uncharacterized | no | HGCF |
| Listeria_phage_LMSP-25 | NC_024360 | P100virus | 2/280 | Uncharacterized | no | HGCF |
| Listeria_phage_LMTA-148 | NC_024787 | P100virus | 2/280 | Uncharacterized | no | HGCF |
| Listeria_phage_vB_LmoM_AG20 | NC_020871 | P100virus | 2/280 | Uncharacterized | no | HGCF |
| Listeria_phage_LP-048 | NC_024359 | P100virus | 2/280 | no | no | HGCF |
| Listeria_phage_LP-083-2 | NC_024383 | P100virus | 2/280 | no | no | HGCF |
| Listeria_phage_LP-125 | NC_021781 | P100virus | 2/280 | no | no | HGCF |
| Listeria_virus_A511 | NC_009811 | P100virus | 2/280 | no | no | HGCF |
| Lactococcus_phage_ul36 | NC_004066 | | 85/277 | no | yes | HGCF |
| Burkholderia_virus_BcepMu | NC_005882 | Bcepmuvirus | 217/222 | yes | no | LGCF |
| Burkholderia_virus_phiE255 | NC_009237 | Bcepmuvirus | 217/222 | Uncharacterized | no | LGCF |
| Salmonella_phage_Fels-1 | NC_010391 | | 218/279 | yes | yes | LGCF |
| Enterobacteria_phage_HK225 | NC_019717 | | 218/279 | yes | yes | Mixed |
| Enterobacteria_phage_mEp237 | NC_019704 | | 218/279 | yes | yes | Mixed |
| Enterobacteria_phage_SfV | NC_003444 | | 83/92 | yes | yes | Mixed |
| Escherichia_virus_N15 | NC_001901 | N15virus | 218/279 | yes | yes | Mixed |
| Salmonella_phage_ST64B | NC_004313 | | 83/92 | yes | yes | Mixed |
| Shigella_phage_SfII | NC_021857 | | 83/92 | yes | yes | Mixed |
| Shigella_phage_SfIV | NC_022749 | | 83/92 | yes | yes | Mixed |
| Phage_Gifsy-1 | NC_010392 | | 218/279 | yes | yes | Unknown |
| Phage_Gifsy-2 | NC_010393 | | 218/279 | yes | yes | Unknown |
| Enterobacteria_phage_HK630 | NC_019723 | Lambdavirus | 13/218 | Uncharacterized | no | Unknown |
| Halorubrum_pleomorphic_virus_1 | NC_012558 | Alphapleolipovirus | 74/134 | Uncharacterized | no | Unknown |
| Idiomarinaceae_phage_1N2-2 | NC_025439 | | 142/191 | Uncharacterized | no | Unknown |
| Marinomonas_phage_P12026 | NC_018269 | | 142/191 | Uncharacterized | no | Unknown |
| Pseudoalteromonas_phage_Pq0 | NC_029100 | | 142/191 | Uncharacterized | no | Unknown |
| Mycobacterium_phage_BigNuz | NC_023692 | Bignuzvirus | 23/143 | yes | yes | Uncharacterized |
| Mycobacterium_phage_Brusacoram | NC_028747 | Fishburnevirus | 23/143 | yes | yes | Uncharacterized |
| Mycobacterium_phage_Donovan | NC_023552 | Fishburnevirus | 23/143 | yes | yes | Uncharacterized |
| Mycobacterium_phage_Fishburne | NC_021302 | Fishburnevirus | 23/143 | yes | yes | Uncharacterized |
| Mycobacterium_phage_Malithi | NC_026605 | Fishburnevirus | 23/143 | yes | yes | Uncharacterized |
| Mycobacterium_phage_Phayonce | NC_028796 | Phayoncevirus | 23/143 | yes | yes | Uncharacterized |
| Mycobacterium_phage_Shipwreck | NC_031261 | | 23/143 | yes | yes | Uncharacterized |
| Gordonia_phage_Twister6 | NC_031052 | Wizardvirus | 139/224 | yes | yes | Uncharacterized |
| Gordonia_phage_Wizard | NC_030913 | Wizardvirus | 139/224 | yes | yes | Uncharacterized |
| Gordonia_phage_Cozz | NC_030941 | | 196/216 | no | no | Uncharacterized |
| Gordonia_phage_Emalyn | NC_031234 | | 196/216 | no | no | Uncharacterized |
| Gordonia_phage_GTE2 | NC_015720 | | 196/216 | no | no | Uncharacterized |
| Halorubrum_pleomorphic_virus_2 | NC_017087 | Alphapleolipovirus | 74/134 | Uncharacterized | no | Uncharacterized |
| Halorubrum_pleomorphic_virus_6 | NC_017089 | Alphapleolipovirus | 74/134 | Uncharacterized | no | Uncharacterized |
| Lactococcus_phage_50101 | NC_031040 | | 85/113 | yes | yes | Uncharacterized |
| Lactococcus_phage_63301 | NC_031017 | | 85/113 | yes | yes | Uncharacterized |
| Salmonella_phage_118970_sal3 | NC_031940 | | 83/92 | Uncharacterized | yes | Uncharacterized |
| Salmonella_phage_118970_sal4 | NC_030919 | | 13/32 | Uncharacterized | yes | Uncharacterized |
| Salmonella_phage_103203_sal5 | NC_031946 | | 13/32 | Uncharacterized | yes | Uncharacterized |
| Enterobacteria_phage_UAB_Phi20 | NC_031019 | | 13/32 | no | yes | Uncharacterized |

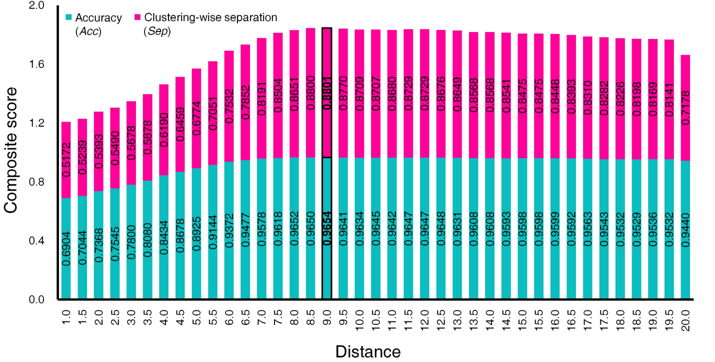

Accuracy (*Acc*)
Clustering-wise separation (*Sep*)
Composite score
Distance
2.0
1.6
1.2
0.8
0.4
0.0
1.0 1.5 2.0 2.5 3.0 3.5 4.0 4.5 5.0 5.5 6.0 6.5 7.0 7.5 8.0 8.5 9.0 9.5 10.0 10.5 11.0 11.5 12.0 12.5 13.0 13.5 14.0 14.5 15.0 15.5 16.0 16.5 17.0 17.5 18.0 18.5 19.0 19.5 20.0
0.5172 0.5239 0.5393 0.5490 0.5678 0.5878 0.6190 0.6459 0.6774 0.7051 0.7532 0.7852 0.8191 0.8504 0.8651 0.8800 0.8801 0.8770 0.8709 0.8707 0.8680 0.8729 0.8729 0.8676 0.8649 0.8568 0.8568 0.8541 0.8475 0.8475 0.8448 0.8393 0.8310 0.8282 0.8226 0.8198 0.8169 0.8141 0.7178
0.6904 0.7044 0.7368 0.7545 0.7800 0.8080 0.8434 0.8678 0.8925 0.9144 0.9372 0.9477 0.9578 0.9618 0.9652 0.9650 0.9654 0.9641 0.9634 0.9645 0.9642 0.9647 0.9647 0.9648 0.9631 0.9608 0.9608 0.9593 0.9598 0.9598 0.9599 0.9592 0.9563 0.9543 0.9532 0.9529 0.9536 0.9532 0.9440

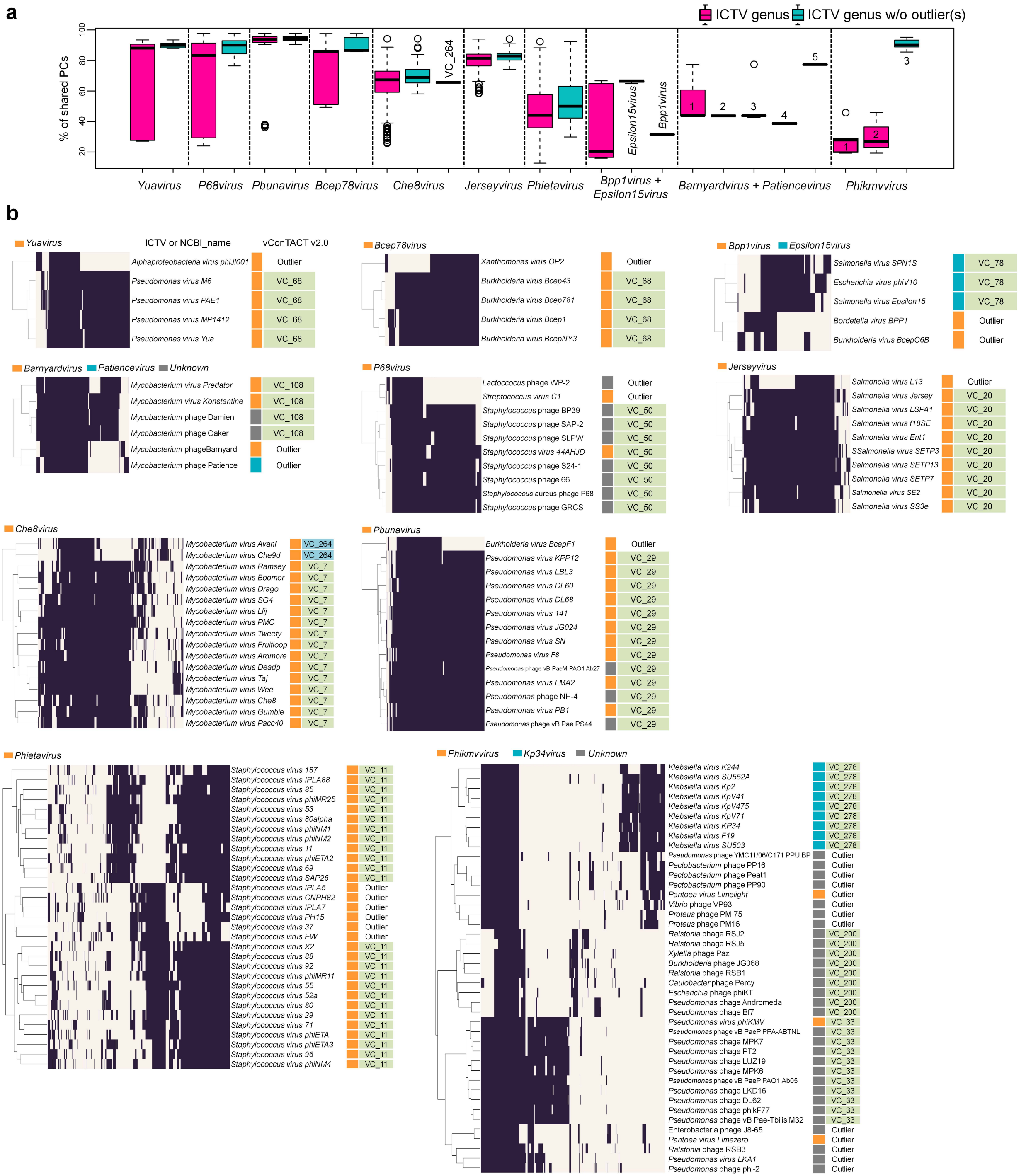

a
ICTV genus
ICTV genus w/o outlier(s)
% of shared PCs
20
40
60
80
100
VC_264
Epsilon15virus
Bpp1virus
1
2
3
4
5
3
1
2
Yuavirus
P68virus
Pbunavirus
Bcep78virus
Che8virus
Jerseyvirus
Phietavirus
Bpp1virus + Epsilon15virus
Barnyardvirus + Patiencevirus
Phikmvvirus
b
Yuavirus
ICTV or NCBI_name
vConTACT v2.0
Alphaproteobacteria virus phiJl001 Outlier
Pseudomonas virus M6 VC_68
Pseudomonas virus PAE1 VC_68
Pseudomonas virus MP1412 VC_68
Pseudomonas virus Yua VC_68
Bcep78virus
Xanthomonas virus OP2 Outlier
Burkholderia virus Bcep43 VC_68
Burkholderia virus Bcep781 VC_68
Burkholderia virus Bcep1 VC_68
Burkholderia virus BcepNY3 VC_68
Bpp1virus
Epsilon15virus
Salmonella virus SPN1S VC_78
Escherichia virus phiV10 VC_78
Salmonella virus Epsilon15 VC_78
Bordetella virus BPP1 Outlier
Burkholderia virus BcepC6B Outlier
Barnyardvirus
Patiencevirus
Unknown
Mycobacterium virus Predator VC_108
Mycobacterium virus Konstantine VC_108
Mycobacterium phage Damien VC_108
Mycobacterium phage Oaker VC_108
Mycobacterium phageBarnyard Outlier
Mycobacterium phage Patience Outlier
P68virus
Lactococcus phage WP-2 Outlier
Streptococcus virus C1 Outlier
Staphylococcus phage BP39 VC_50
Staphylococcus phage SAP-2 VC_50
Staphylococcus phage SLPW VC_50
Staphylococcus virus 44AHJD VC_50
Staphylococcus phage S24-1 VC_50
Staphylococcus phage 66 VC_50
Staphylococcus aureus phage P68 VC_50
Staphylococcus phage GRCS VC_50
Jerseyvirus
Salmonella virus L13 Outlier
Salmonella virus Jersey VC_20
Salmonella virus LSPA1 VC_20
Salmonella virus f18SE VC_20
Salmonella virus Ent1 VC_20
SSalmonella virus SETP3 VC_20
Salmonella virus SETP13 VC_20
Salmonella virus SETP7 VC_20
Salmonella virus SE2 VC_20
Salmonella virus SS3e VC_20
Che8virus
Mycobacterium virus Avani VC_264
Mycobacterium virus Che9d VC_264
Mycobacterium virus Ramsey VC_7
Mycobacterium virus Boomer VC_7
Mycobacterium virus Drago VC_7
Mycobacterium virus SG4 VC_7
Mycobacterium virus Llij VC_7
Mycobacterium virus PMC VC_7
Mycobacterium virus Tweety VC_7
Mycobacterium virus Fruitloop VC_7
Mycobacterium virus Ardmore VC_7
Mycobacterium virus Deadp VC_7
Mycobacterium virus Taj VC_7
Mycobacterium virus Wee VC_7
Mycobacterium virus Che8 VC_7
Mycobacterium virus Gumbie VC_7
Mycobacterium virus Pacc40 VC_7
Pbunavirus
Burkholderia virus BcepF1 Outlier
Pseudomonas virus KPP12 VC_29
Pseudomonas virus LBL3 VC_29
Pseudomonas virus DL60 VC_29
Pseudomonas virus DL68 VC_29
Pseudomonas virus 141 VC_29
Pseudomonas virus JG024 VC_29
Pseudomonas virus SN VC_29
Pseudomonas virus F8 VC_29
Pseudomonas phage vB PaeM PAO1 Ab27 VC_29
Pseudomonas virus LMA2 VC_29
Pseudomonas phage NH-4 VC_29
Pseudomonas virus PB1 VC_29
Pseudomonas phage vB Pae PS44 VC_29
Phietavirus
Staphylococcus virus 187 VC_11
Staphylococcus virus IPLA88 VC_11
Staphylococcus virus 85 VC_11
Staphylococcus virus phiMR25 VC_11
Staphylococcus virus 53 VC_11
Staphylococcus virus 80alpha VC_11
Staphylococcus virus phiNM1 VC_11
Staphylococcus virus phiNM2 VC_11
Staphylococcus virus 11 VC_11
Staphylococcus virus phiETA2 VC_11
Staphylococcus virus 69 VC_11
Staphylococcus virus SAP26 VC_11
Staphylococcus virus IPLA5 Outlier
Staphylococcus virus CNPH82 Outlier
Staphylococcus virus IPLA7 Outlier
Staphylococcus virus PH15 Outlier
Staphylococcus virus 37 Outlier
Staphylococcus virus EW Outlier
Staphylococcus virus X2 VC_11
Staphylococcus virus 88 VC_11
Staphylococcus virus 92 VC_11
Staphylococcus virus phiMR11 VC_11
Staphylococcus virus 55 VC_11
Staphylococcus virus 52a VC_11
Staphylococcus virus 80 VC_11
Staphylococcus virus 29 VC_11
Staphylococcus virus 71 VC_11
Staphylococcus virus phiETA VC_11
Staphylococcus virus phiETA3 VC_11
Staphylococcus virus 96 VC_11
Staphylococcus virus phiNM4 VC_11
Phikmvvirus
Kp34virus
Unknown
Klebsiella virus K244 VC_278
Klebsiella virus SU552A VC_278
Klebsiella virus Kp2 VC_278
Klebsiella virus KpV41 VC_278
Klebsiella virus KpV475 VC_278
Klebsiella virus KpV71 VC_278
Klebsiella virus KP34 VC_278
Klebsiella virus F19 VC_278
Klebsiella virus SU503 VC_278
Pseudomonas phage YMC11/06/C171 PPU BP Outlier
Pectobacterium phage PP16 Outlier
Pectobacterium phage Peat1 Outlier
Pectobacterium phage PP90 Outlier
Pantoea virus Limelight Outlier
Vibrio phage VP93 Outlier
Proteus phage PM 75 Outlier
Proteus phage PM16 Outlier
Ralstonia phage RSJ2 VC_200
Ralstonia phage RSJ5 VC_200
Xylella phage Paz VC_200
Burkholderia phage JG068 VC_200
Ralstonia phage RSB1 VC_200
Caulobacter phage Percy VC_200
Escherichia phage phiKT VC_200
Pseudomonas phage Andromeda VC_200
Pseudomonas phage Bf7 VC_200
Pseudomonas virus phiKMV VC_33
Pseudomonas phage vB PaeP PPA-ABTNL VC_33
Pseudomonas phage MPK7 VC_33
Pseudomonas phage PT2 VC_33
Pseudomonas phage LUZ19 VC_33
Pseudomonas phage MPK6 VC_33
Pseudomonas phage vB PaeP PAO1 Ab05 VC_33
Pseudomonas phage LKD16 VC_33
Pseudomonas phage DL62 VC_33
Pseudomonas phage phiKF77 VC_33
Pseudomonas phage vB Pae-TbilisiM32 VC_33
Enterobacteria phage J8-65 Outlier
Pantoea virus Limezero Outlier
Ralstonia phage RSB3 Outlier
Pseudomonas virus LKA1 Outlier
Pseudomonas phage phi-2 Outlier

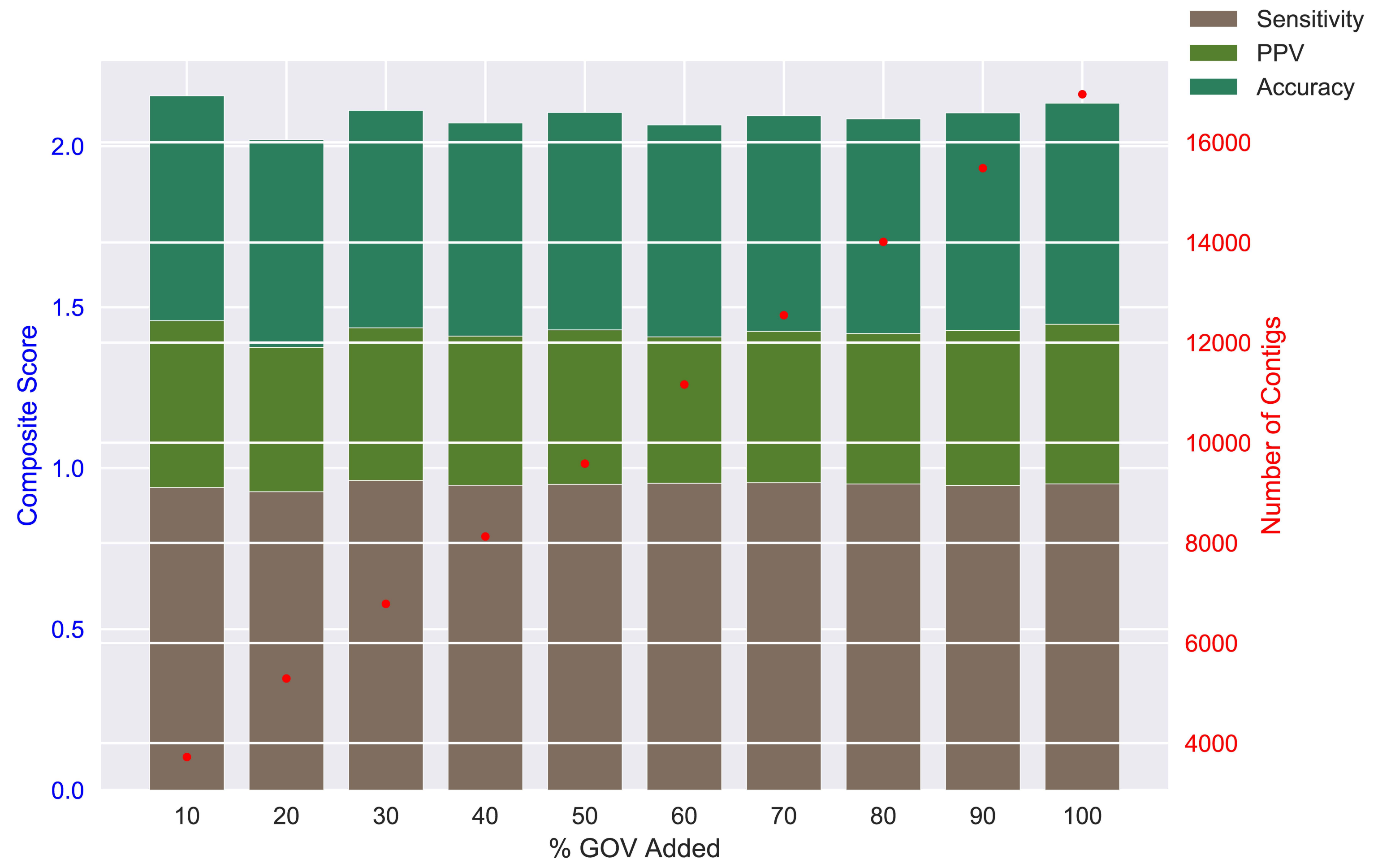

Sensitivity
PPV
Accuracy
Composite Score
0.0
0.5
1.0
1.5
2.0
Number of Contigs
4000
6000
8000
10000
12000
14000
16000
10
20
30
40
50
60
70
80
90
100
% GOV Added

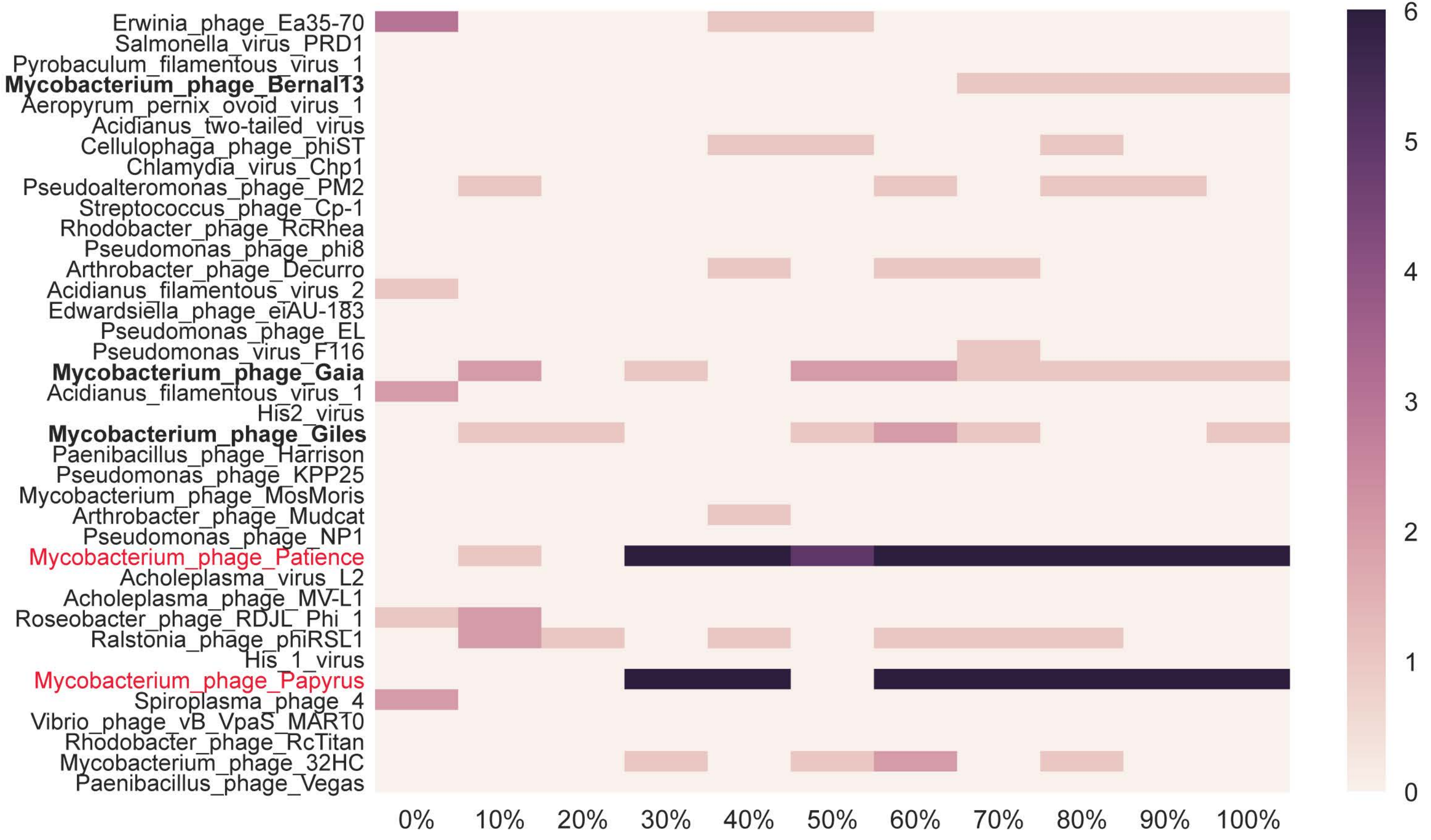
Erwinia_phage_Ea35-70
Salmonella_virus_PRD1
Pyrobaculum_filamentous_virus_1
Mycobacterium_phage_Bernal13
Aeropyrum_pernix_ovoid_virus_1
Acidianus_two-tailed_virus
Cellulophaga_phage_phiST
Chlamydia_virus_Chp1
Pseudoalteromonas_phage_PM2
Streptococcus_phage_Cp-1
Rhodobacter_phage_RcRhea
Pseudomonas_phage_phi8
Arthrobacter_phage_Decurro
Acidianus_filamentous_virus_2
Edwardsiella_phage_eiAU-183
Pseudomonas_phage_EL
Pseudomonas_virus_F116
Mycobacterium_phage_Gaia
Acidianus_filamentous_virus_1
His2_virus
Mycobacterium_phage_Giles
Paenibacillus_phage_Harrison
Pseudomonas_phage_KPP25
Mycobacterium_phage_MosMoris
Arthrobacter_phage_Mudcat
Pseudomonas_phage_NP1
Mycobacterium_phage_Patience
Acholeplasma_virus_L2
Acholeplasma_phage_MV-L1
Roseobacter_phage_RDJL_Phi_1
Ralstonia_phage_phiRSL1
His_1_virus
Mycobacterium_phage_Papyrus
Spiroplasma_phage_4
Vibrio_phage_vB_VpaS_MAR10
Rhodobacter_phage_RcTitan
Mycobacterium_phage_32HC
Paenibacillus_phage_Vegas
0%
10%
20%
30%
40%
50%
60%
70%
80%
90%
100%
6
5
4
3
2
1
0

| Inflation factor | ICCC | Number of clusters |
|---|---|---|
| 1.0 | 0.603587766 | 52 |
| 1.2 | 0.603587766 | 52 |
| 1.4 | 0.64600304 | 141 |
| 1.6 | 0.642135592 | 188 |
| 1.8 | 0.618643803 | 214 |
| 2.0 | 0.615766124 | 237 |
| 2.2 | 0.61545278 | 243 |
| 2.4 | 0.609649475 | 250 |
| 2.6 | 0.601978682 | 256 |
| 2.8 | 0.605250475 | 262 |
| 3.0 | 0.599840822 | 267 |
| 3.2 | 0.596050936 | 271 |
| 3.4 | 0.596473778 | 275 |
| 3.6 | 0.592123251 | 277 |
| 3.8 | 0.590415616 | 281 |
| 4.0 | 0.594927805 | 284 |
| 4.2 | 0.60501266 | 286 |
| 4.4 | 0.603078366 | 288 |
| 4.6 | 0.600483223 | 289 |
| 4.8 | 0.601237419 | 291 |
| 5.0 | 0.595581164 | 294 |

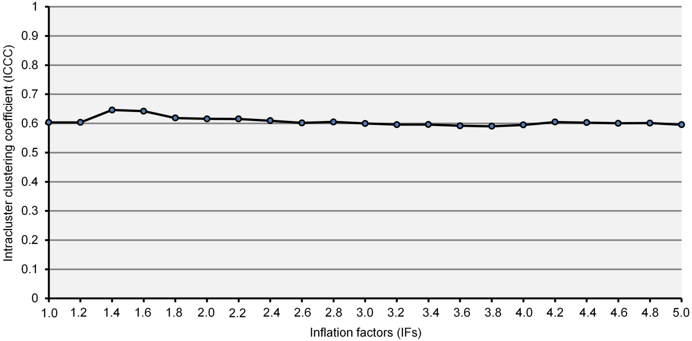
